# Estimating SNP heritability in presence of population substructure in biobank-scale datasets

**DOI:** 10.1101/2020.08.05.236901

**Authors:** Zhaotong Lin, Souvik Seal, Saonli Basu

## Abstract

SNP heritability of a trait is measured by the proportion of total variance explained by the additive effects of genome-wide single nucleotide polymorphisms (SNPs). Linear mixed models are routinely used to estimate SNP heritability for many complex traits. The basic concept behind this approach is to model genetic contribution as a random effect, where the variance of this genetic contribution attributes to the heritability of the trait. This linear mixed model approach requires estimation of ‘relatedness’ among individuals in the sample, which is usually captured by estimating a genetic relationship matrix (GRM). Heritability is estimated by the restricted maximum likelihood (REML) or method of moments (MOM) approaches, and this estimation relies heavily on the GRM computed from the genetic data on individuals. Presence of population substructure in the data could significantly impact the GRM estimation and may introduce bias in heritability estimation. The common practice of accounting for such population substructure is to adjust for the top few principal components of the GRM as covariates in the linear mixed model. Here we propose an alternative way of estimating heritability in multi-ethnic studies. Our proposed approach is a MOM estimator derived from the Haseman-Elston regression and gives an asymptotically unbiased estimate of heritability in presence of population stratification. It introduces adjustments for the population stratification in a second-order estimating equation and allows for the total phenotypic variance vary by ethnicity. We study the performance of different MOM and REML approaches in presence of population stratification through extensive simulation studies. We estimate the heritability of height, weight and other anthropometric traits in the UK Biobank cohort to investigate the impact of subtle population substructure on SNP heritability estimation.

## 1 Introduction

Fundamental to the study of the inheritance is the partitioning of the total phenotypic variation into genetic and environmental components (Visscher et al., 2008). Using twin studies, the phenotypic variance can be partitioned to include the variance of an additive genetic effect, shared and non-shared environmental effects. The ratio of the genetic variance component to the total phenotypic variance is the proportion of genetically controlled variation and is termed as the ‘narrow-sense heritability’. As shown in the recent review of more than 17,000 twin studies (Polderman et al., 2015), heritability provides useful information to estimate familial recurrence risk of disease, to inform about the genetic architecture of the trait, and to generate an upper bound for disease risk prediction.

In recent years, the genome-wide association studies (GWAS) are gaining momentum with the availability of whole genome sequencing data. Heritability is routinely being estimated from the genome-wide data on variants (single nucleotide polymorphisms or SNPs), which is often termed as ‘SNP heritability’. Traditionally, SNP heritability is estimated by fitting variance components models with restricted maximum likelihood (REML) approach. These approaches partition the phenotypic covariance matrix of all individuals into a genetic similarity matrix and a random variation matrix (Yang et al., 2010; Lee et al., 2011, 2012; Ripke et al., 2013; AR et al., 2014; Locke et al., 2015).

However, with the large sample size, for example, biobanks that assay hundreds of thousands of individuals (UK Biobank (Biobank, 2014), Precision Medicine cohort (Ashley, 2015), Millions Veterans Program (Gaziano et al., 2016)), existing heritability estimation methods such as REML-based methods become computationally expensive and memory intensive, and thus can be difficult to apply.

Alternatively, there are method-of-moments (MOM) estimators for heritability. LD-score regression approach (Bulik-Sullivan et al., 2015) estimates heritability by regressing the summary statistics from single variant association analysis in a GWAS on linkage disequilibrium (LD) scores. A version of Haseman-Elston approach (Haseman, 1972) for heritability estimation provides a method of moments estimator for the heritability parameter by associating phenotypic covariance values with genetic covariance estimates. There are several recent work on extending these Method of Moments estimators (Ge et al., 2015; Schwartzman et al., 2019; Ma and Dicker, 2019; Hou et al., 2019) to make it more computationally feasible for large sample sizes and more robust to linkage disequilibrium.

Presence of population substructure can significantly bias the heritability estimation (Browning and Browning, 2011). Confounding can occur because of not accounting for the phenotypic differences among different sub-populations due to differences in environmental influences. Moreover, population substructure introduces differences in allele frequencies across sub-populations. Current heritability estimation methods primarily work well in samples from a homogeneous population. However, diverse populations are increasingly being used to conduct GWAS to improve fine-mapping of relevant variants. Recently, Conomos et al. (2016) performed an association study in the admixed Hispanic Community Health Study/Study of Latinos (HCHS/SOL) samples, where many biomedical traits in HCHS/SOL displayed heterogeneous variances across ethnic groups. Modeling this heteroscedasticity reduced genomic inflation. Conomos et al. (2016) estimated the underlying ethnic groups through multi-dimensional scaling and estimated distinct ethnic clusters to implement such correction. It is often desirable to implement such corrections on a continuous scale, for example, modeling heterogeneity in variances along the axes of genetic variation.

In this paper, we propose a strategy to correct for the impact of population stratification on heritability estimation with Haseman-Elston regression. Our approach does not require classifying individuals into discrete sub-populations, rather the corrections are implemented as a function of axes of genetic variation. Another huge advantage of our proposed approach is that it is a method of moments estimator and can provide computationally efficient estimates of heritability even for large biobank-scale datasets.

The rest of the paper is arranged as follows. We describe few existing approaches to estimate heritability. We propose our modified Haseman-Elston estimator and show the equivalency with the heritability estimator proposed by Ge et al. (2015). We further demonstrate that this estimator gives an unbiased estimate of heritability in presence of 2 discrete sub-populations. We explore the performance of the estimator under various alternative models and compare the performance with existing approaches. Finally we estimate heritability for a number of anthropometric traits on UK Biobank dataset.

## 2 Methods

Linear mixed models are emerging as the method of choice for association testing in genome-wide association studies (GWAS) because they account for both population stratification and cryptic relatedness and achieve increased statistical power by jointly modeling all genotyped markers.

### 2.1 Existing Approaches

Here we first introduce a general mixed-effect model to quantify how genes influence phenotypes. Suppose the data consists of *P* SNPs on *N* subjects. For a subject *i* (*i* = 1, …, *N*), **y**_*i*_ is a normally distributed continuous outcome, **C**_*i*_ is the vector of covariates, ***β*** is a vector of fixed effects, **Z**_*i*_ is a *P* × 1 vector of genetic variants from a GWAS. The outcome **y**_*i*_ depends on **Z**_*i*_ through the following mixed effect model, **y** = **C*β*** + **Zu** + ***ϵ***, with 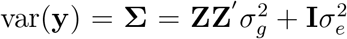, where **u** is a vector of SNP effects with 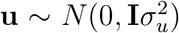, **I** is an *N* × *N* identity matrix, and ***ϵ*** is a vector of residual effects with 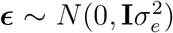. **Z** is a standardized *N* × *P* genotype matrix with the *is*-th element 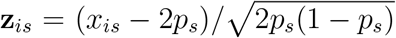, where *x*_*is*_ is the number of copies of the reference allele for the *s*-th SNP of the *i*-th individual and *p*_*s*_ is the frequency of the reference allele. If we define **A** = **ZZ**′/*P* and define 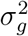 as the variance explained by all the SNPs, i.e, 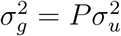, with *P* being the number of SNPs, then the above linear model reduces to

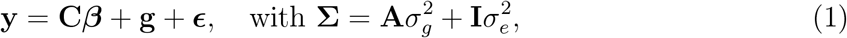

where **g** is an *N* × 1 vector of the total genetic effects of the individuals with 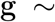 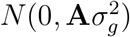, and **A** is interpreted as the genetic relationship matrix (GRM) between individuals. Note that the genetic relatedness **A**_*ij*_ between the *i*-th individual and the *j*-th individual is measured by the dot product of their standardized genotypes and then divided by the number of markers, 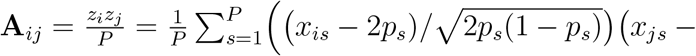 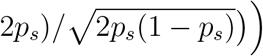. The heritability *h*^2^ of the trait **y** is defined as 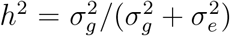.

We are interested in estimating the parameters 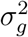 and 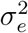.

#### Maximum Likelihood Estimation

We assume 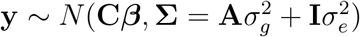. The likelihood of the data is given by:

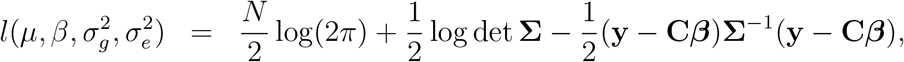

The estimation of the mixed effect model mentioned above is performed through maximum likelihood estimation. The software GCTA (Yang et al., 2011) uses the iterative restricted maximum likelihood (REML) algorithm to estimate the variance components 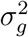 and 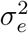 in the model 2 and gives an estimate of heritability by 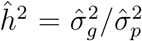, where 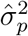 is the estimated phenotypic total variance 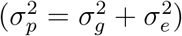.

However, this mixed-model methods can easily become computationally intractable as the sample size increases. Recently, there have been attempts to generate computationally scalable algorithms to implement this mixed models on biobank scale data (Loh et al., 2015). However, these approaches still encounter computational challenges on large biobanks. Moreover, even subtle population substructure could significantly impact the heritability estimation (Conomos et al., 2016).

#### Method of Moments Approach

The method of moments (Haseman-Elston regression (Haseman, 1972), LDscore regression (Bulik-Sullivan et al., 2015), MMHE (Ge et al., 2017)) approaches are another set of widely used methods for estimating heritability *h*^2^ under Equation 1. We will next provide short overview of these different approaches.

##### Haseman-Elston (HE) Regression

Generally we assume that the GRM is normalized with its diagonal entries all equal 1 and **y** is centered and that Equation 1 holds. One of the classical moment estimators for *h*^2^ comes from the least squares regression coefficient for regressing **y**_*i*_**y**_*j*_ on **A**_*ij*_ for all *i* < *j*. This is because Equation 1 implies that *E*(**y**_*i*_**y**_*j*_|**A**) = *h*^2^**A**_*ij*_ for *i* ≠ *j*. The heritability can be estimated from the following equation:

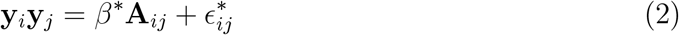

Note *β*^*^ = *h*^2^ is the heritability parameter. The corresponding estimator for *h*^2^ is 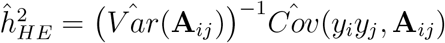. Note that

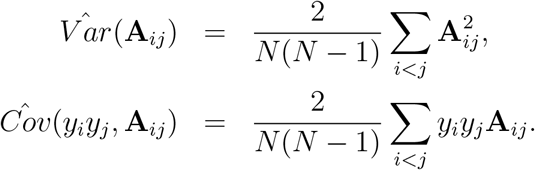

Henderson (1984) used least squares in this way to estimate 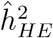. This approach is also referred to as Haseman-Elston (HE) regression (Haseman, 1972).

A modification to the above approach is to consider two estimating equations for both parameters 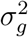 and 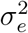:

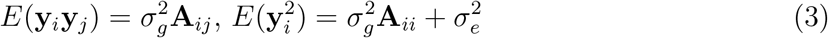

Again, least square estimation can be used to produce unbiased estimates of 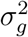 and 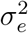.

##### Linkage Disequilibrium Score Regression

As an alternative method, linkage disequilibrium score (LDSC) regression (Bulik-Sullivan et al., 2015) has become a popular approach for estimating SNP heritability from summary statistics. LDSC estimates SNP heritability by regressing squared per-SNP univariate regression test statistics on corresponding “LD Scores”, defined as estimates of the sum of squared correlations for a given SNP with all other SNPs within a region. The main advantage of this approach is that it can utilize the summary statistics generated from a GWAS, which are publicly available. The asymptotic equivalence between LDSC approach and the HE regression has been derived under certain assumptions (Chen, 2014; Bulik-Sullivan, 2015). However, while an effective and computationally efficient approach, LDSC relies on a number of assumptions, including independence of individuals to compute the summary statistics, binning of LD scores. This introduces some arbitrariness to consider the approach analytically and limit the assessment of its theoretical properties.

##### MMHE

Ge et al. (2016, 2017) proposed this MOM estimator, which is closely related to Haseman-Elston regression estimator (Haseman, 1972) and is equivalent to LD score regression estimator under certain situations (Bulik-Sullivan et al., 2015; Bulik-Sullivan, 2015). Specifically, in the presence of covariates, i.e., **y** = **C*β*** + **g** + ***ϵ***, an *N* × (*N* − *k*) matrix **U** always exists, such that **U**^*T*^ **U** = **I**, **UU**^*T*^ = **H**, **U**^*T*^ **C** = **0** and **H** = **I** − **C**(**C**^*T*^ **C**)^−1^**C**^*T*^. Applying **U**^*T*^ to both sides of the model gives **U**^*T*^ **y** = **U**^*T*^ **g** + **U**^*T*^ **ϵ**. The covariance structure of the transformed trait is 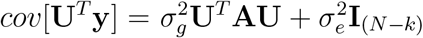. Then by converting a matrix into a vector by stacking its columns, an ordinary least squares (OLS) estimator of 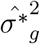, 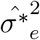 can be obtained by solving the linear system:

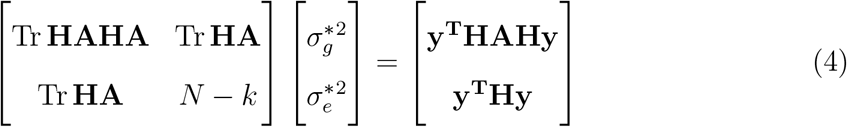

and obtain

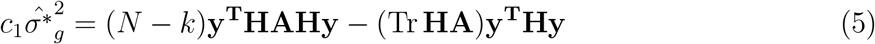

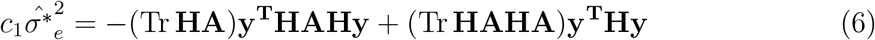

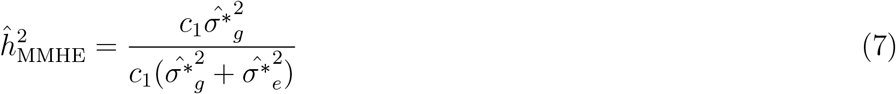

where **H** = **I** − **P** and **P** = **C**(**C^T^C**)^−**1**^**C^T^**

### 2.2 Proposed Adjusted HE method

The MOM or the likelihood-based approaches generally assumes a homogeneous population. In a sample of diverse ancestry, these existing methods could produced very biased estimates of heritability. The proposed concept is motivated by the idea is that population substructure causes differences in allele frequencies as well as differences in trait distributions among the sub-populations. Not accounting for such differences could significantly introduce bias in the estimation of heritability. The standard approach is to adjust for principal components (PCs) estimated from the GRM **A** as covariates in Equation 1. The MOM-based approaches provide an useful alternative for large samples, but it is always not clear how such adjustments for substructure could be implemented in the approach. In this paper, we propose a two-step strategy to adjust for population substructure in estimating heritability using Haseman-Elston regression (Haseman, 1972). We first perform a regression on the mean level of the trait:

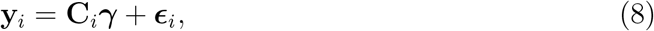

by regressing out covariates **C**, which might consist of *k* PCs and other covariates such as sex and age. We assume that the residuals **y**′ still preserve the same information and structure of heritability, i.e., 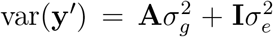. For the second step, we consider two different approaches to account for population stratification. One approach is to adjust the allele frequencies with PCs and recompute GRM **A** as implemented in PC-Relate (Conomos et al., 2016), and then apply the standard HE regression to obtain heritability estimate (referred as ‘PC-Relate-HE’). Another novel way is that we introduce PC-based corrections in Equation 3. We will refer to the method as Adjusted-HE approach. The potential difference between PC-Relate-HE and Adjusted-HE is that, the former only adjusts the GRM entries for population substructure, whereas Adjusted-HE introduces correction to both GRM entries and to the total variance of outcome.

#### HE regression with PC-Relate adjusted GRM (PC-Relate-HE)

PC-Relate (Conomos et al., 2016) is a PCA-based method for robust estimation of IBD-sharing probabilities and kinship coefficients that is applicable to general samples with population structure. Consider the linear regression 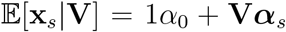 where **x**_*s*_ is the vector of genotype values for all samples at SNP *s* and **V** is a matrix whose columns correspond to the top *k* PCs from PC-Air (Conomos et al., 2015). The fitted values from this regression can be used as prediction of individual-specific allele frequencies from the PCs: 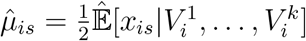. Then the PC-Relate estimator of the genetic relationship coefficient **A**_*ij*_ for individual *i* and *j* is

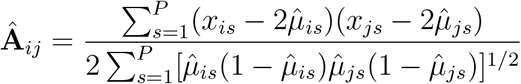

where 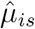 and 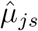 are the estimated individual-specific genotype mean for individual *i* and *j*, respectively, at SNP *s*. Then we estimate the heritability with this PC-adjusted GRM with the standard HE regression.

##### Unstandardized-Adjusted-HE (UAdj-HE)

In this method, we consider the following estimating equation:

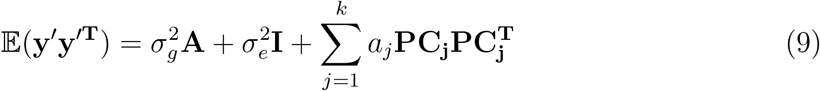

where **y**′ = (**I** − **C**(**C^T^C**)^−**1**^**C**^**T**^)**y** is the residual of the regression in Equation 8.

One could use ordinary least square (OLS) approach to estimate 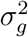 and 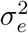 here. We have also derived a closed form estimator of SNP heritability using the following equations (see Appendix A),

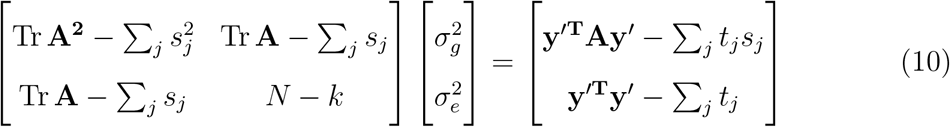

and obtain

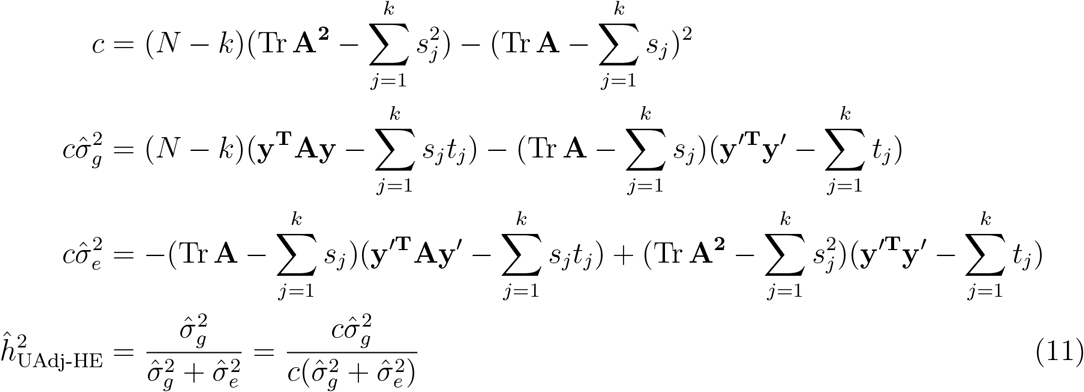

where 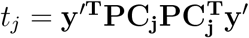, 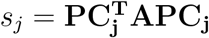.

##### Standardized-Adjusted-HE (SAdj-HE)

If we have the residuals **y**′ standardized by the sample mean and variance of Equation 8, then based on Equation 2 we have the following estimating equation:

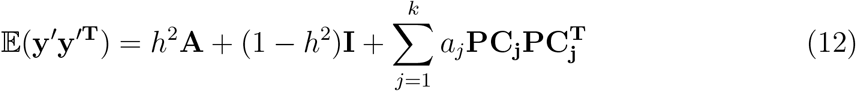

where **y**′ is the standardized residual of the Equation 8.

Then we can obtain the estimate (derivation is shown in Appendix A)

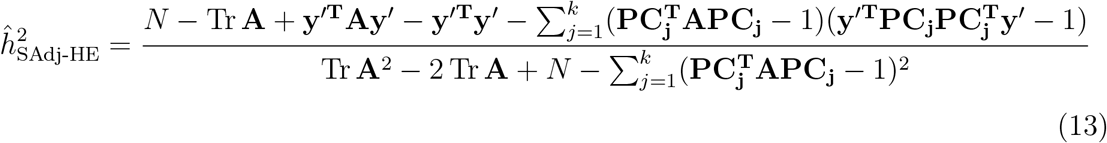

We have shown that, in the presence of two distinct sub-populations, this estimator with the first PC product adjustment can give us unbiased estimate (See Appendix B). To calculate the variance of Adj-HE estimators, we can make two similar assumptions as Ge et al. (2017) :(1) the off-diagonal elements in the empirical GRM matrix **A** are small and the diagonal elements are close to 1, such that **A** ≈ **I** and (2) the phenotypic variance can be estimated precisely. Therefore, we have 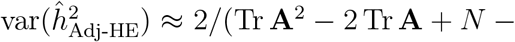 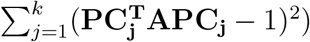. And with the assumption of independence among samples, we use the standard error of the OLS estimator derived from Equation 12 and Equation 9.

#### 2.2.1 Relationship between MMHE and Adjusted-HE

Our proposed UAdj-HE is equivalent to the MMHE approach (Ge et al., 2017), when we adjust for the PCs in Equation 9 and Equation 4 are the same PCs computed from the entire GRM **A**. Assume the set of covariates, **C** only consists of the PCs, i.e., 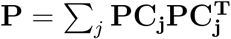,

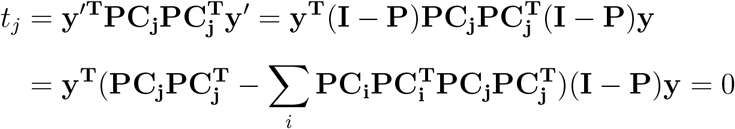

Then Equation 10 reduces to

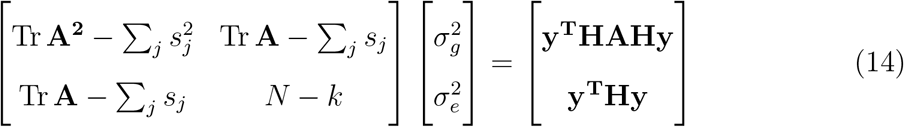

where 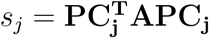. Moreover, in Equation 4,

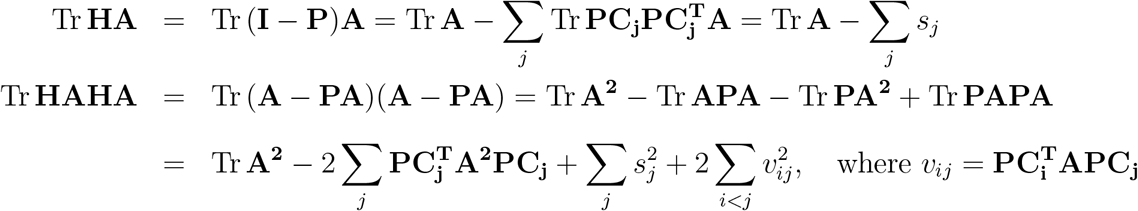

The only difference between Equation 4 and Equation 14 is Tr **HAHA** and 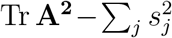. In the case of using the set of **PC**s calculated from the same GRM matrix **A**, we can use the fact that **APC_j_** = λ_*j*_**PC_j_**, then *v*_*ij*_ = 0 and 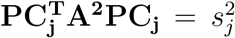. As a result, 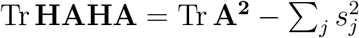 and two methods are equivalent.

However if the set of covariates contain other covariates such as age, sex or if the PCs are estimated by sampling an independent subset of markers from the given set of markers, the two methods are not equivalent and may produce different estimates for heritability. However, unless there is significant impact of other covariates on the variance and covariance of the trait, we do not expect the estimates to differ significantly.

## 3 Results

We conducted extensive simulation studies and real data analysis to evaluate the performance of Adjusted-HE, MMHE, PC-Relate-HE and GCTA-REML methods with and without principal components adjustment.

### 3.1 Simulation Studies

We considered 3 different simulation setup to assess the performance of these methods in presence of allele frequency differences and differences in trait distributions among the sub-populations. We simulated allele frequencies of 15,000 SNPs for each distinct population using the Balding-Nichols model (Balding and Nichols, 1995). The SNPs were assumed to be uncorrelated. For each SNP *s*, the allele frequency *p*_0*s*_ in the ancestral population was drawn from a uniform distribution on [0.1, 0.9]. In simulation 1 and 2, for each sub-population *k*, the allele frequency *p*_*ks*_ was generated from a beta distribution with parameters *p*_0*s*_(1 − *θ*_*k*_)/*θ*_*k*_ and (1 − *p*_0*s*_)(1 − *θ*_*k*_)/*θ*_*k*_. The parameter *θ*_*k*_ was set to a common value in simulation 1 and we varied *θ*_*k*_ across sub-populations in simulation 2. In simulation 3, the allele frequency of *k*-th population *p*_*ks*_ at SNP *s* was generated from a beta distribution with parameters *p*_(*k*−1)*s*_(1 −*θ*_*k*_)/*θ*_*k*_ and (1 −*p*_(*k*−1)*s*_)(1 −*θ*_*k*_)/*θ*_*k*_, where *θ*_*k*_ was set to a common value (*k* = 1, 2, 3, 4 and *s* = 1, 2, …, 15000). Next, we simulated the genotypes **x**_*ks*_ of individuals in sub-population *k* from a binomial distribution *Bin*(2, *p*_*ks*_) assuming Hardy-Weinberg equilibrium. We only considered SNPs with MAF > 0.05 and selected *m* (15,000 × *p*_*causal*_) SNPs as causal variants with an effect size 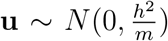, where *p*_*causal*_ was the proportion of causal variants. Then the residual effects *e*_*k*_ were generated from a normal distribution with mean of 0 and variance of 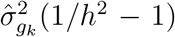, where 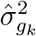 is the empirical variance of **X**_*k*_**u**, **X**_*k*_ is a *n*_*k*_ × *m* unstandardized causal genotype matrix, **u** is a *m* × 1 vector of causal effects and *h*^2^ is the given heritability. Finally, we simulated phenotype *y*_*ki*_ of individual *i* in population *k* as *y*_*ki*_ = **X**_*ki*_**u** + *e*_*ki*_ + *a*_*k*_, where **X**_*ki*_ is a 1 × *m* vector of causal SNPs of individual *i* in sub-population *k* and *a*_*k*_ is a population-intercept to make the means of sub-populations more different.

We considered 4 discrete populations, each with 1,000 samples, and *p*_*causal*_ = 0.02 in all simulations. The results were based on 100 replications for each setup. We used SAdj-HE, UAdj-HE, MMHE, PC-Relate-HE and GCTA-REML with no PC adjustment to 5 PCs adjustments for estimating heritability.

#### Simulation 1

We considered *θ*_*k*_ of 0.01 (closely related sub-populations) and 0.1 (more divergent populations) with *h*^2^ set to 0.8. We also considered scenarios without or with population-intercept (i.e., (*a*_1_, *a*_2_, *a*_3_, *a*_4_) = (0, 0, 0, 0) or (*a*_1_, *a*_2_, *a*_3_, *a*_4_) = (0, 1, 2, 3)). Figure 1 shows the results of scenario when *θ*_*k*_ = 0.1 and *h*^2^ = 0.8. The density curves of MMHE and UAdj-HE were identical, since they are equivalent methods as demonstrated in Section 2.2.1. As expected, the heritability estimation for MOM approaches stabilized after adjusting for 3 PCs as the sample had 4 different sub-populations. When the mean differences in **y** across sub-populations were small (*a*_*k*_ = 0), GCTA-REML handled the impact of population substructure well, even when there were no PC adjustments (Figure 1 top panel). But when population means were different, i.e., (*a*_1_, *a*_2_, *a*_3_, *a*_4_) = (0, 1, 2, 3), GCTA-REML showed bias in heritability estimation when less than 3 PCs were used as covariates (Figure 1 bottom panel).

**Figure 1.**
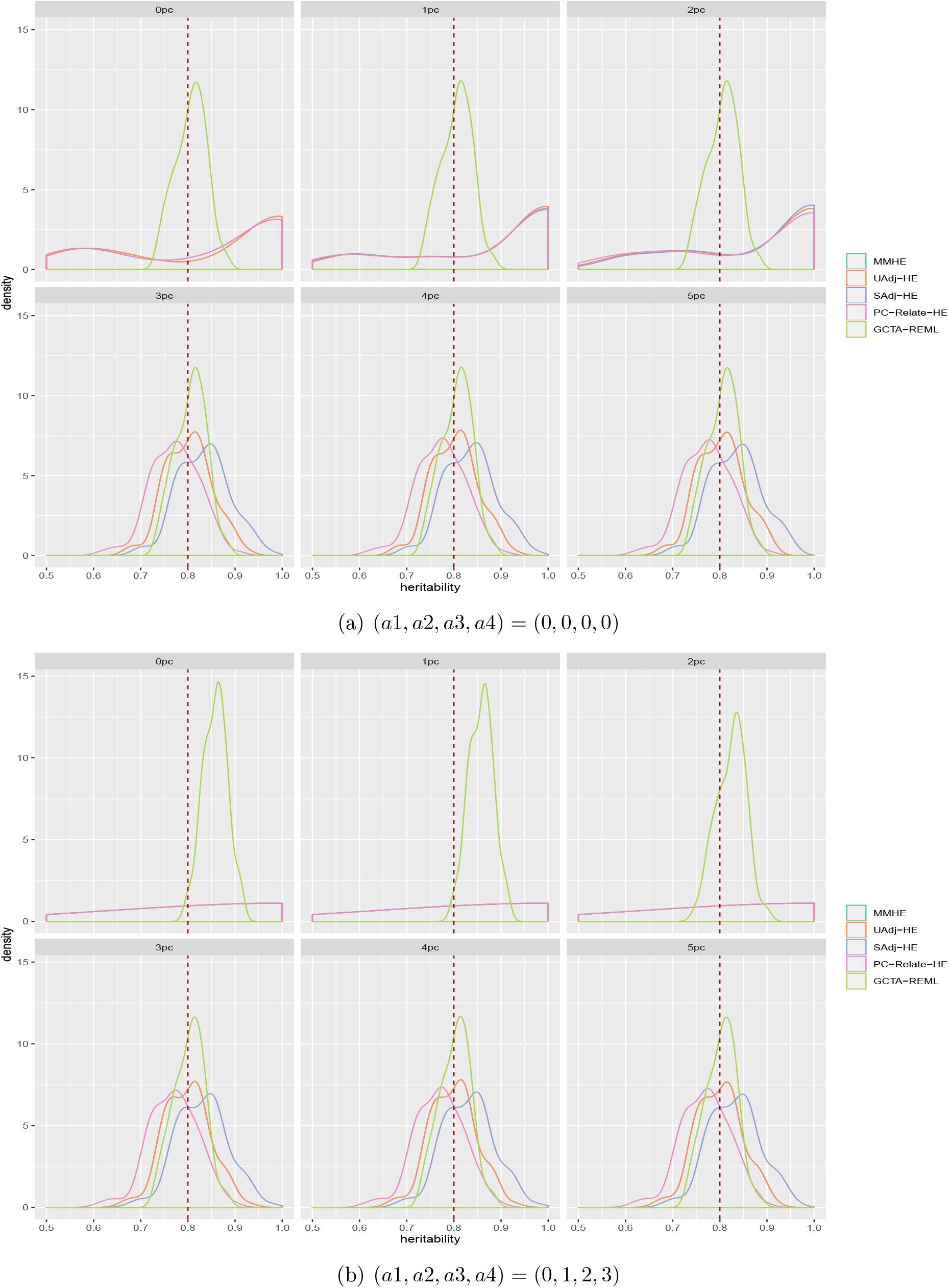
Simulation 1: Heritability estimation of different methods with 0 to 5 PCs adjustment with *h*^2^ = 0.8 and *θ*_*k*_ = 0.1

Figure 2 shows boxplots of heritability estimates over 100 replicates for different methods. We show results for all the methods adjusted for 3 PCs. When 4 populations were similar (*θ*_*k*_ = 0.01 and *a*_*k*_ = 0), all methods performed well while GCTA-REML had the smallest variance. But when the populations were genetically similar (*θ*_*k*_ = 0.01) but with different population intercept (*a*_*k*_ ≠ 0), heritability was underestimated by all methods. This is possibly due to the fact that the PCs were not informative to distinguish between the sub-populations and hence PC adjustments could not account for the differences among the subpopulations. When the populations were more diverse (*θ*_*k*_ = 0.1), PC-Relate-HE showed downward bias, whereas GCTA-REML, MMHE and Adj-HE estimates were biased upward.

**Figure 2.**
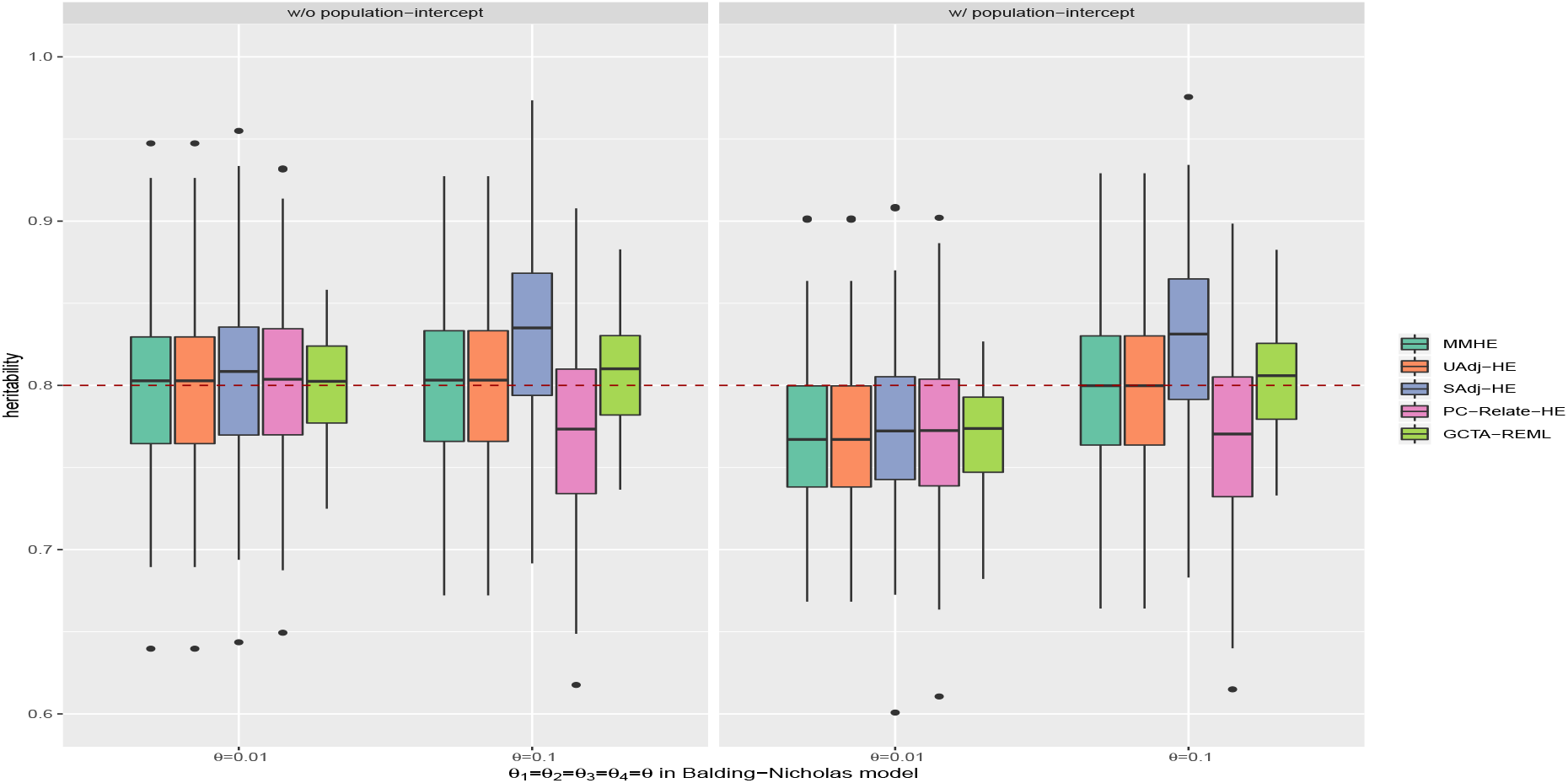
Simulation 1: Heritability estimation of different methods with 3 PCs adjustment across 100 replicates. Dashed line is the true *h*^2^ = 0.8. Left: *a*_*k*_ = 0. Right: (*a*_1_, *a*_2_, *a*_3_, *a*_4_) = (0, 1, 2, 3)

#### Simulation 2

In this simulation setup, we varied *θ*_*k*_ to consider different pairwise similarities between each of the sub-populations with the ancestral population. The parameters *θ*_1_, *θ*_2_, *θ*_3_, *θ*_4_ were set to 0.05, 0.1, 0.15 and 0.2 respectively. As a result, the variances of **y** in different sub-populations were different compared to simulation 1 with a common *θ*_*k*_ (Supplementary Figure S1). Figure 3 shows the result of scenarios without population-intercept *a*_*k*_ (left panel) and with population-intercept *a*_*k*_ (right panel). The methods MMHE, UAdj-HE and GCTA-REML showed marginal overestimation, but PC-Relate-HE significantly underestimated heritability.

**Figure 3.**
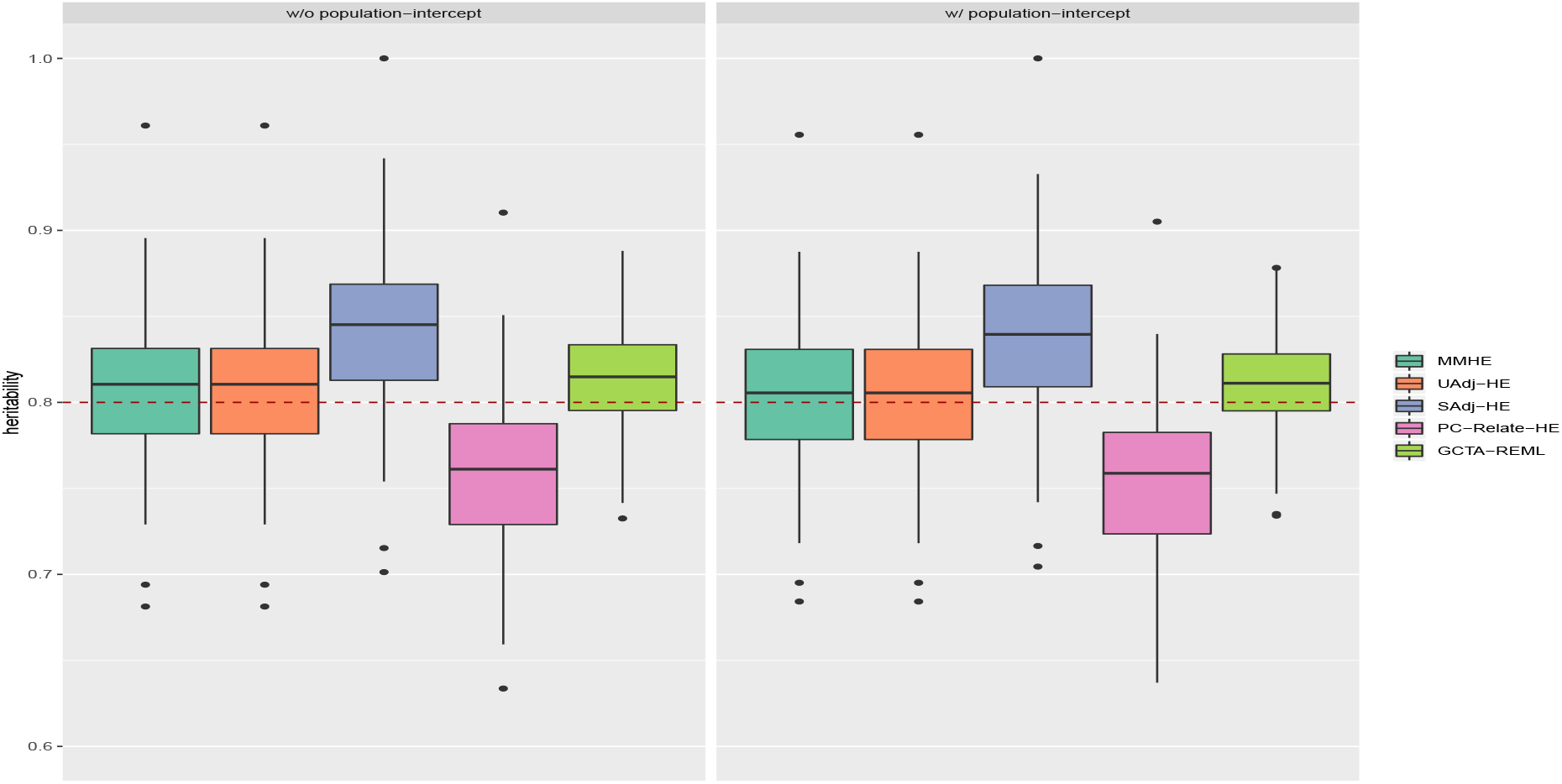
Simulation 2: Heritability estimation of different methods with 3 PCs adjustment with *h*^2^ = 0.8 and *θ*_1_, *θ*_2_, *θ*_3_, *θ*_4_ = (0.05, 0.1, 0.15, 0.2). Left: *a*_*k*_ = 0; Right: (*a*_1_, *a*_2_, *a*_3_, *a*_4_) = (0, 1, 2, 3)

#### Simulation 3

In this simulation, we used *θ*_*k*_s to represent genetic similarity between population pair (*k* − 1, *k*), *k* = 2, 3, 4. This simulation generated more diverse sub-populations as compared to Simulation 1 and Simulation 2. Similar to simulation 2, the variances of **y** in each sub-population were more different as *θ*_*k*_ increasing (Supplementary Figure S2). We estimated the pairwise Fst value between sub-populations using the empirical Bayes estimator in FinePop package (Kitada et al., 2007) (Supplementary Table S1). As we increased *θ*_*k*_, PC-Relate-HE showed increasing downward bias and GCTA-REML and SAdj-HE biased upward more. The approaches while UAdj-HE and MMHE performed the best with smallest bias among the methods (Figure 4).

**Figure 4.**
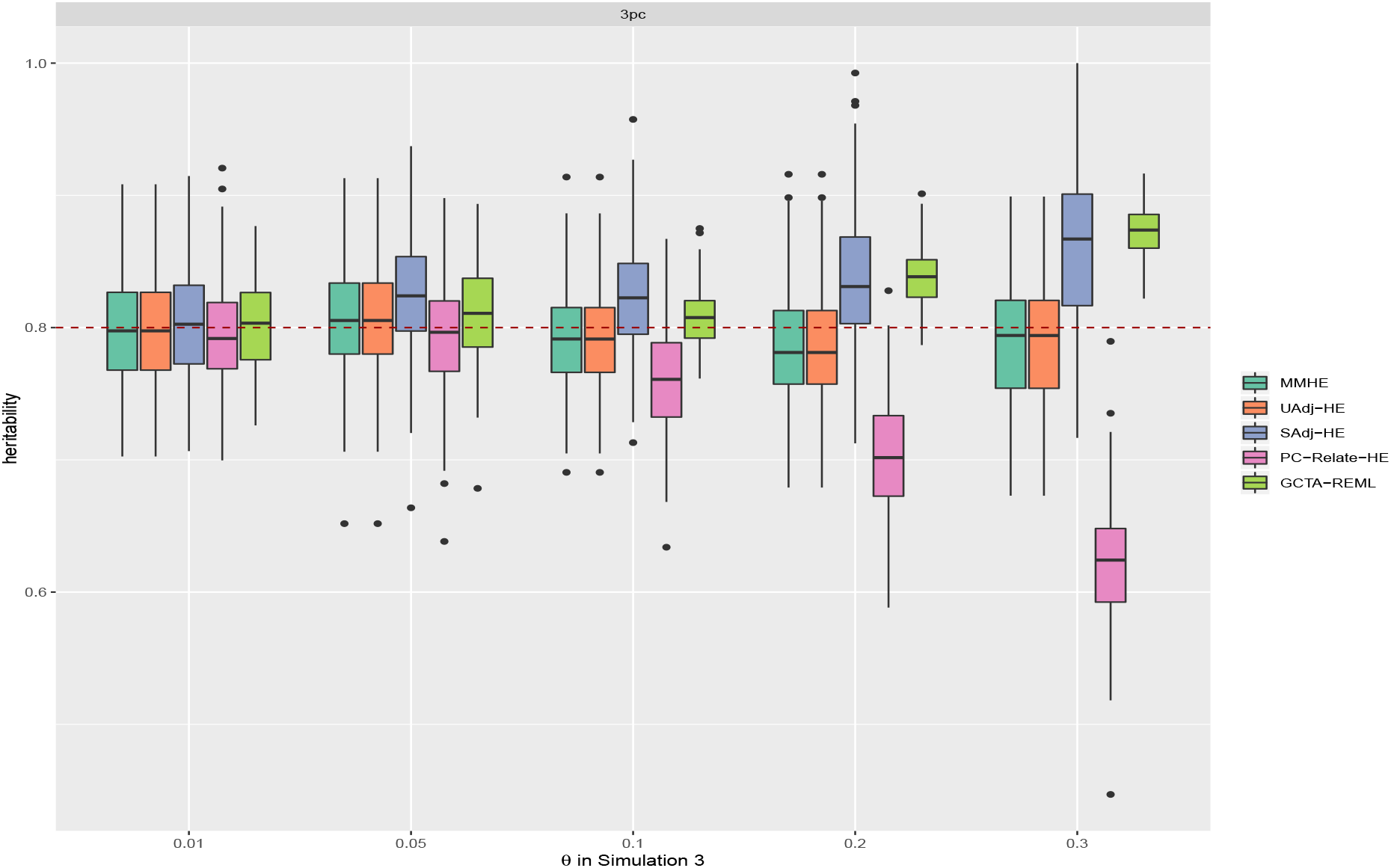
Simulation 3: Heritability estimation of different methods with 3 PCs adjustment with *h*^2^ = 0.8 and *θ*_1_ = *θ*_2_ = *θ*_3_ = *θ*_4_ = *θ*, *a*_*k*_ = 0.

In general, in the presence of population substructure, all the MOM estimators performed better than the original HE regression without any PC adjustment, when adjusted for sufficient PCs. PC-Relate-HE showed underestimation, especially when sub-populations were more diverse. Our proposed approach and REML showed slight overestimation in most scenarios. When the variances of phenotype differed across sub-populations, PC-Relate-HE underestimated severely. This is probably due to fact that PC-Relate-HE only adjusted the GRM for population substructure, whereas Adj-HE approaches corrected both the GRM and the total variance of **y** for population substructure and showed little bias in the estimation of heritability. As expected, GCTA-REML estimates had the smallest variance compared to the other MOM estimators, since the data was simulated from a normal distribution, however, it suffered from overestimation when there was heterogeneity of variances across sub-populations. SAdj-HE showed more bias in estimation over UAdj-HE, probably because of the inaccuracy in standardization of **y**, especially when there was heterogeneity of variances across sub-populations. In terms of averaged computational time for each replicate of simulation, PC-Relate took about 1700 seconds to run PC-Air and create the PCs-adjusted GRM for 4,000 samples, whereas it only took less than 30 seconds totally to calculate the PCs and construct the standard GRM in PLINK and GCTA; and MMHE and GCTA-REML took about 3 seconds and 20 seconds in heritability estimation respectively whereas Adjusted-HE only took less than a second running on Haswell E5-2680v3 processors.

### 3.2 Real Data Analysis: UK Biobank

The UK Biobank data has approximately 800,000 markers and comprises 488,377 samples. We leveraged the QC information released by UK Biobank (Bycroft et al., 2018) and used SNPs that passed all QC tests in 106 batches. We removed the samples that had mismatch between inferred sex and self-reported sex, samples that were identified as outliers in heterozygosity and missing rates, and samples that were in the kinship table (Biobank, 2015). We further excluded SNPs that had high missing rate (>1.5%), low minor allele frequency (<1%) and subjects that had high missing genotype rate (>1%). 305,639 samples and 566,647 markers remained for the following analysis after QC. It is also worth mentioning that we did not restrict our analysis to subjects that were self-reported white British. The majority of the samples was British, but there were people with other ethnicities (Supplementary Table S3, S4). PCs were computed after LD pruning and removing long-range LD-regions (Abdellaoui et al., 2013). We pruned the SNPs after removing long-range LD-regions so that pairwise *r*^2^ < 0.2 among the remaining markers for windows of 1000 markers and a step-size of 80 markers. We computed PCs using the pruned data with 247,135 markers.

We studied the performance of different approaches on a subset of the 305,639 samples, since GCTA-REML cannot be handle this large sample size. We sampled 45,510 subjects from 305,639 subjects who are self-reported White, Asian or Black, and applied GCTA-REML, LD score regression, MMHE, SAdj-HE and UAdj-HE on this sub-sample. We analyzed 7 quantitative phenotypes including height, weight, BMI, waist, levels of the diisobutyl phthalate (DiBP), systolic blood pressure (SysBP), hip and waist circumference. We adjusted for the top few PCs and other covariates such as age and height as recommended in Ge et al. (2017), except for UK Biobank assessment center (Table 1). For the Adjusted-HE methods, we first regressed out PCs and other covariates from each phenotype, then applied the closed-form formula or performed least-square estimation using Equation 12 to estimate heritability. For the MMHE method, we considered PCs and other covariates in the matrix **P**. We adjusted for covariates and PCs when conducting GWAS for LDscore regression, and for GCTA-REML, we included PCs and other covariates in Equation 1 while performing the REML estimation.

**Table 1.** A comparison of different methods of estimating heritability for 45k samples with 10 PCs to correct population substructure

|  | Height<sup>1</sup> | Weight<sup>2</sup> | BMI<sup>1</sup> | DiBP<sup>1</sup> | sysBP<sup>1</sup> | Hip<sup>3</sup> | Waist<sup>3</sup> |
| --- | --- | --- | --- | --- | --- | --- | --- |
| LDSC | 0.523 | 0.196 | 0.197 | 0.119 | 0.136 | 0.099 | 0.047 |
| GCTA-REML | 0.648 | 0.346 | 0.344 | 0.180 | 0.196 | 0.142 | 0.165 |
| MMHE | 0.685 | 0.294 | 0.289 | 0.150 | 0.158 | 0.121 | 0.150 |
| UAdj-HE | 0.638 | 0.271 | 0.269 | 0.140 | 0.147 | 0.112 | 0.138 |
| SAdj-HE | 0.646 | 0.273 | 0.271 | 0.140 | 0.148 | 0.112 | 0.139 |

<sup>1</sup> Sex, age are included in other covariates.
<sup>2</sup> Sex, age and height are included in other covariates.
<sup>3</sup> Sex, age, height and weight are included in other covariates.

#### 3.2.1 45k sample

45,510 subjects were sampled from 305,639 subjects who are self-reported White, Asian or Black (Supplementary Table S2). Similar to our simulation above, when all SNPs were used to compute PCs, UAdj-HE gave the same result as MMHE even when we adjusted for age and sex (results not shown here), which might indicate that other covariates such as sex and age did not affect the variance and covariance of the traits significantly. Table 1 shows the heritability estimation of different methods when 10 PCs (computed based on pruned SNPs) and appropriate covariates were adjusted. We can see that, Adjusted-HE estimates were slightly lower than MMHE, if PCs were not computed based on all SNPs that were used to generate the empirical GRM. However, Adj-HE methods were computationally more efficient (Table 3). LDSC regression method produced substantially smaller estimates compared to other approaches, which indicates severe underestimation by LDSC approach in presence of population stratification. GCTA-REML gave the highest estimates of heritability in most of the cases (except for height), which might also be overestimation due to heterogeneity of variances as shown in our simulation study. Also, as expected, in a large sample size, UAdj-HE and SAdj-HE gave similar results.

#### 3.2.2 305k sample

As we mentioned before, this 305k UK Biobank cohort is a collection of samples from different ethnic backgrounds including White (British, Irish), Mixed (White and Black Caribbean, White and Black African, White and Asian), Asian (Indian, Pakistani, Bangladeshi, Chinese, Asian British) and Black (Caribbean, African, Black British) (Supplementary Figure S3). The estimated variability for different traits across ethnicity is shown in Supplementary Table S4, and it shows different outcome variances across sub-populations. We applied Adjusted-HE corrected for 10 PCs and other covariates on this cohort (Table 2). Compared to the Adjusted-HE results in the 45k sample, most of the estimations increased except for weight and BMI. The results also demonstrated a good consistency with previous results (Hou et al., 2019; Ge et al., 2017) and our approach was computationally very efficient. The MMHE approach needs to take block-columns GRM as input, which is not the standard GRM format provided by GCTA. The GRM file generated by GCTA only stores the lower triangular and diagonal entries of the GRM. In contrast, our proposed method can take the standard GCTA file with whole GRM as input in a more efficient way and use formula (13) if the machine has sufficient memory, otherwise, it can also read part GRM files generated by GCTA and conducts the adjusted regression in parallel. The analysis for 10 PCs adjustment only took several minutes when jobs were paralleled on Haswell E5-2680v3 processors (Table 3).

**Table 2.** Heritability estimation of 305k samples with Adj-HE corrected for PCs and other covariates

| Method | Height | Weight | BMI | DiBP | sysBP | Hip | Waist |
| --- | --- | --- | --- | --- | --- | --- | --- |
| SAdj-HE + 10 PCs | 0.689 | 0.271 | 0.273 | 0.169 | 0.156 | 0.119 | 0.160 |
| UAdj-HE + 10 PCs | 0.684 | 0.271 | 0.272 | 0.169 | 0.155 | 0.119 | 0.159 |
| Ge et al. (2017)<sup>1</sup> | 0.685 | 0.277 | 0.274 | 0.184 | 0.156 | 0.106 | 0.155 |

<sup>1</sup> 108,158 self-reported white-British and 486,175 SNPs were used.

**Table 3.**
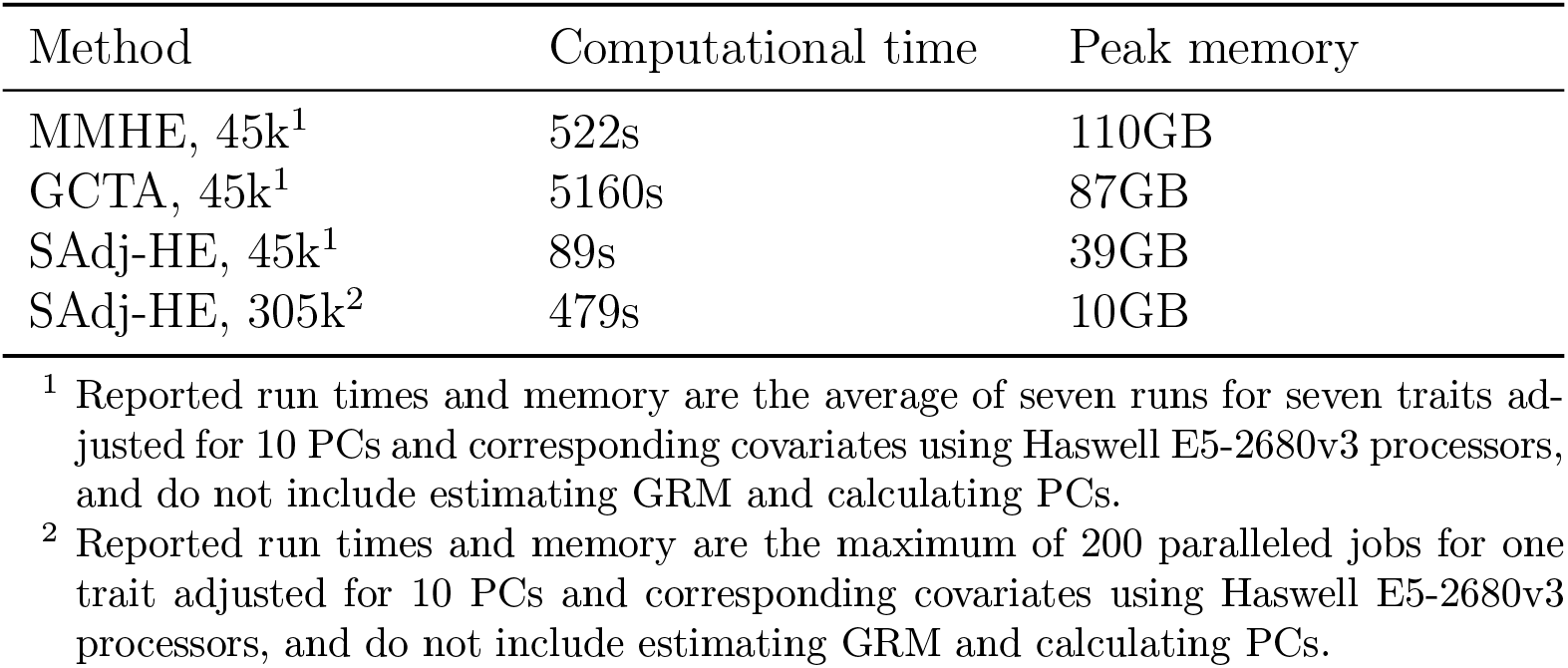
Computational performance of MMHE, GCTA-REML and Adjusted-HE

| Method | Computational time | Peak memory |
| --- | --- | --- |
| MMHE, 45k<sup>1</sup> | 522s | 110GB |
| GCTA, 45k<sup>1</sup> | 5160s | 87GB |
| SAdj-HE, 45k<sup>1</sup> | 89s | 39GB |
| SAdj-HE, 305k<sup>2</sup> | 479s | 10GB |

<sup>1</sup> Reported run times and memory are the average of seven runs for seven traits adjusted for 10 PCs and corresponding covariates using Haswell E5-2680v3 processors, and do not include estimating GRM and calculating PCs.
<sup>2</sup> Reported run times and memory are the maximum of 200 paralleled jobs for one trait adjusted for 10 PCs and corresponding covariates using Haswell E5-2680v3 processors, and do not include estimating GRM and calculating PCs.

## 4 Discussion

SNP-heritability, the proportion of variation in the phenotype attributable to the additive effects of a given set of SNPs, is a fundamental quantity in genetics and provides an upper bound to the risk explained by genetic prediction models. Traditionally, SNP-heritability is estimated by fitting variance components models with restricted maximum likelihood (REML). But these REML-based methods are not scalable to large biobank data. An alternative method is Haseman-Elston regression which is a moment-based method and is computationally much more efficient for large-scale datasets. In recent years, more GWAS are conducted on diverse population and the presence of population substructure can bias SNP-heritability estimation severely. For example, the difference in the outcome variance by ethnicity will impact the estimation. Another major impact of ethnicity is that the genetic relationship matrix **A** will be wrongly computed due to the difference in allele frequencies by ethnicity. Principal components estimated from GRM are usually used to account for population structure. A classical way to adjust for PCs in REML-based methods is to include them as fixed effects in the mixed linear model; and PCs can also be adjusted in the estimation of GRM using methods such as PC-Relate (Conomos et al., 2015). However, it is still unclear how to incorporate such corrections in different existing moment based approaches for estimating heritability.

In this paper, we proposed a computationally efficient MOM estimator of SNP-heritability in presence of population substructure, which can be easily applied on large scale biobank data. We have derived the estimator from the classical Haseman-Elston regression by adding product terms of PCs and have shown the equivalence with MMHE under specific conditions. We have also demonstrated the unbiasedness of our proposed estimator in presence of two discrete sub-populations. Another flexibility of our proposed approach is that it would be relatively easy to allow the heritability differ by ethnicity. One could incorporate multiple interaction terms in the Adj-HE approach (Equation 12 and Equation 9) to allow for such ethnic differences.

We conducted a number of simulations to study the performance of Adjusted-HE and other methods including GCTA-REML, MMHE and PC-Relate-HE for heritability estimation. In simulation studies under a variety of population substructure configurations, we showed that if not adjusting for PCs, MOM estimators are biased severely in the presence of population substructure; and the estimates stabilized after adjusting for PCs. When sub-populations had similar outcome mean and variances, GCTA-REML estimates were stable even without any PC adjustment, but it also needed PC adjustments to stabilize if outcome means or variances were different. We also noticed increasing downward bias for PC-Relate-HE and increasing upward bias for GCTA-REML, when the subpopulations were increasingly different in outcome variances. The UAdj-HE approach always maintained the smallest bias in these scenarios.

In the real data application on UK Biobank, we analyzed 7 quantitative traits including height, weight, BMI, systolic blood pressure, diastolic blood pressure, waist circumference and hip circumference. We compared Adjusted-HE to other widely used methods including GCTA-REML and LDSC regression on a small subset of 45k individuals, and also applied Adjusted-HE on a full sample of 305k individuals in a computationally efficient way. The results showed that LDSC regression tended to give underestimation. GCTA-REML gave higher heritability estimation than our methods for all traits, likely due to difference in trait variances by ethnicity. Our Adj-HE estimates were generally close to the estimates reported for anthropometric traits from other studies (Yang et al., 2015).

Despite the computational efficiency, our proposed method has a few limitations. Our two-step Adj-HE approach assumes that there is no impact of other covariates such as age and sex on the variance of the outcome. It also slightly underestimated heritability when the PCs were computed using a subset of markers in the GRM. Moreover, it is a bit unclear in terms of how many PC adjustments would be necessary to capture the impact of population stratification.

## Appendix A Derivation of Adjusted-HE formula

## UAdj-HE

For unstandardized **y**′ (only mean-centered), we have

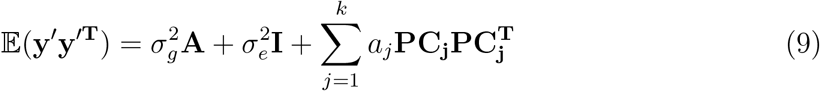

where **y**′ is the residual after the regression step.

Using the method of moment and the Frobenius matrix norm to solve it,

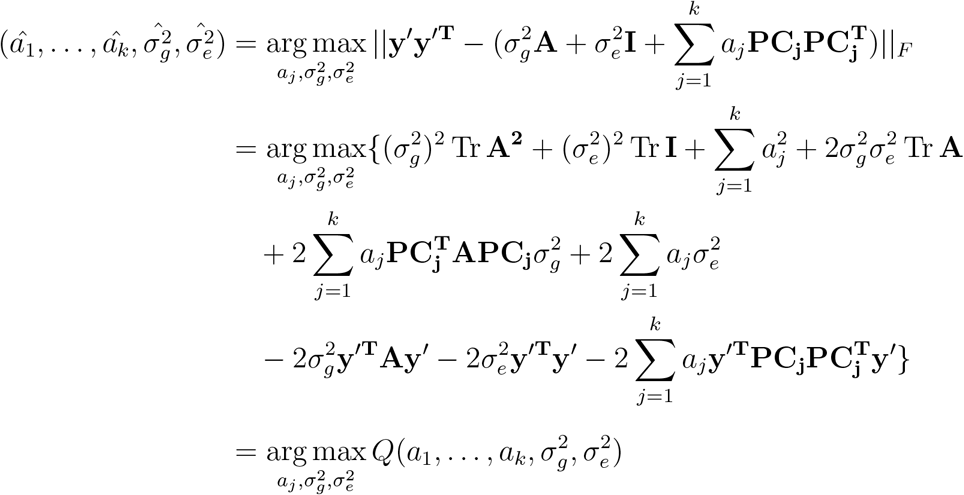

Let 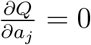, 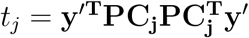, 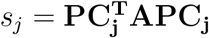, we have 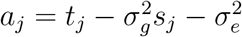.

Let 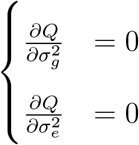, we have

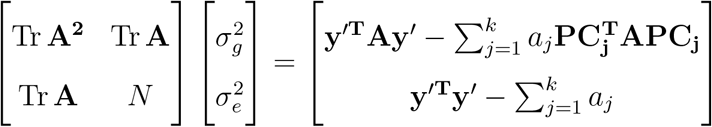

Plug in *a*_*j*_, it becomes

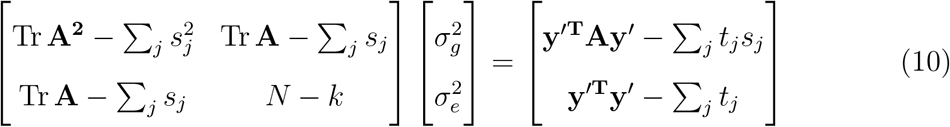

Then

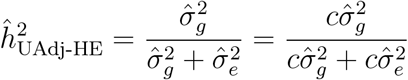

where

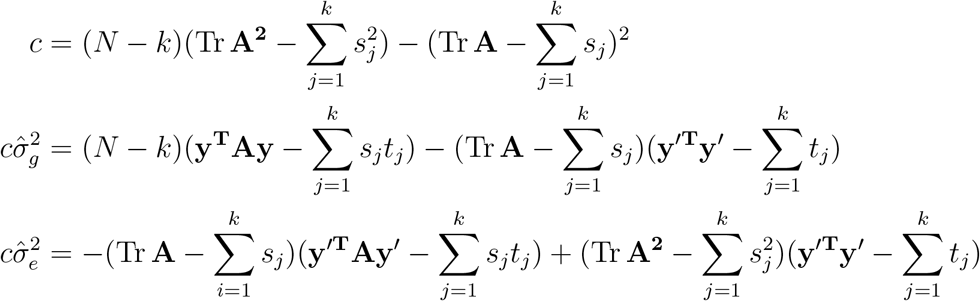

## SAdj-HE

For standardized **y**′ (in both mean and variance), we have

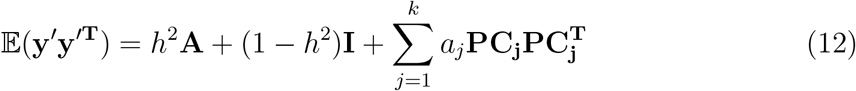

where **y**′ is the standardized residual after first step.

Using the same idea to solve Equation 12, we have

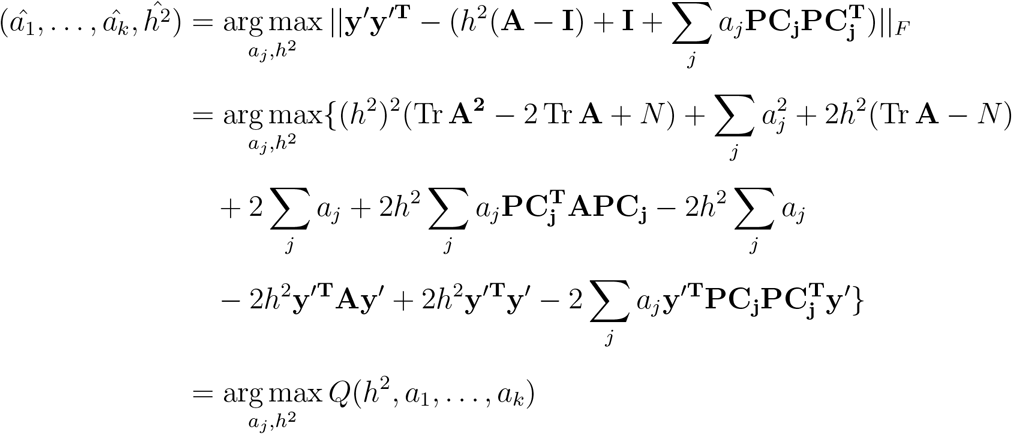

Let 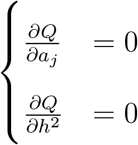, we have

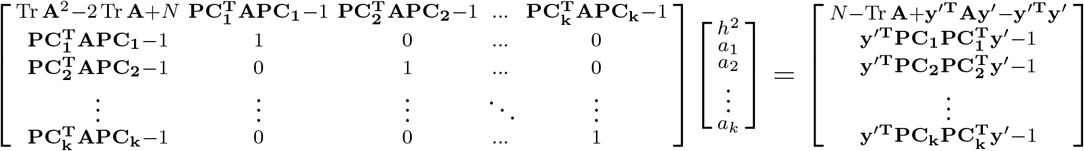

Then we can obtain

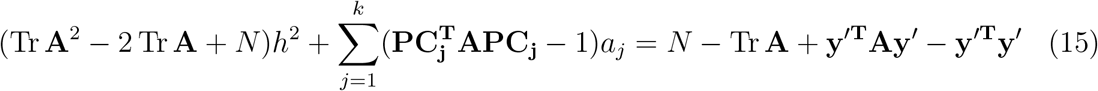

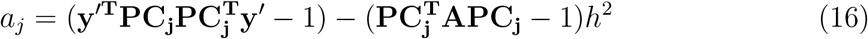

Substitute (16) into (15) we have

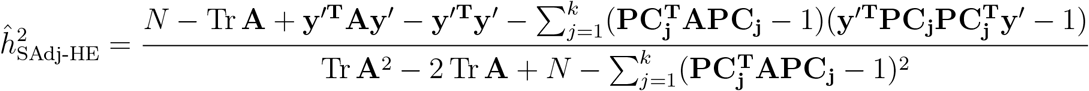

## Appendix B Expectation of the heritability estimates

Assuming that different clusters affect allele frequencies (GRM) only and not the mean directly, and the variance of **y** is estimated precisely by the sample variance. When we only consider PCs from the full GRM in the adjustment, formula 13 can be written as

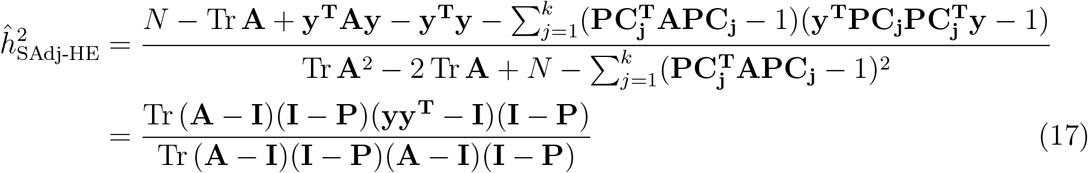

We have, *E*(**yy**^*T*^) = *h*^2^**A**_*True*_ + (1 − *h*^2^)**I** where **A**_*True*_ is the true GRM (where the standardisation of the genotype matrix has been done based on subclusters). Since, we do not know the subclusters, we do not know **A**_*True*_ either. Denote the GRM **A** which we generally work with as **A**_*usual*_ (where the genotype matrix is standardised overall). also denote **H** = **I** − **P**. Under these notations the expression of heritability in Equation 17 above can be written as,

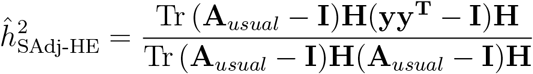

and its expectation can be obtained as,

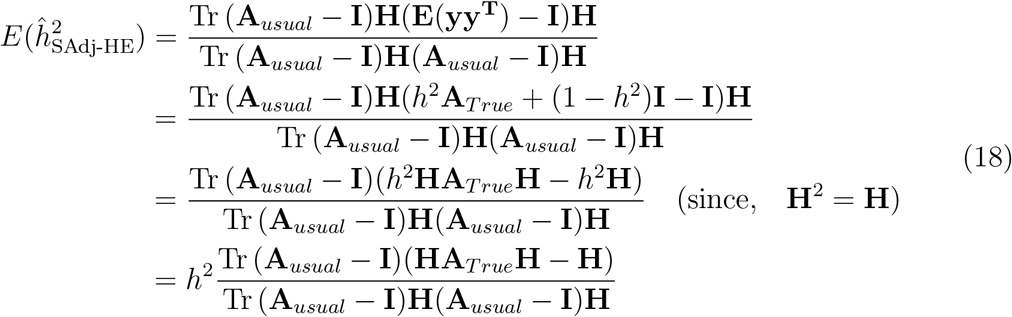

## B.1 Theory with clusters

Suppose there are 2 clusters of individuals, i.e., *k* = 1, 2. Let the allele frequencies of *s*-th SNP for *i*-th individual from cluster *k* be *p*_*ks*_. The total number of individuals is *N* = *n*_1_ + *n*_2_. Let *p*_*s*_ = *r*_1_*p*_1*s*_ + *r*_2_*p*_2*s*_ with 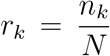. We make an assumption that 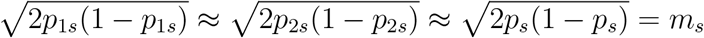 (this is a reasonable assumption since even if *p*_1*s*_, *p*_2*s*_ are much different 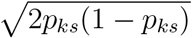’s are not, see Figure 5).

**Figure 5.**
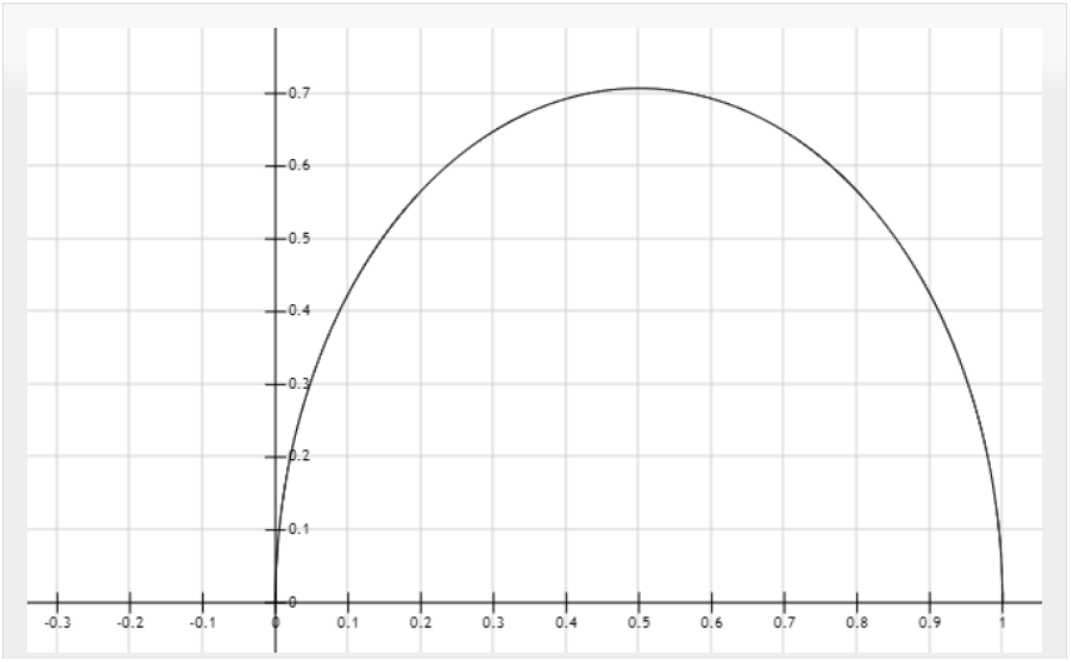
See how the variance function behaves for varying *p*, the function 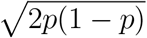 takes pretty close values even when *p* takes highly different values

The corresponding raw(unscaled) genotype matrix be 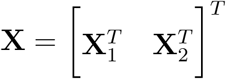 where **X**_*k*_ is of dimension *n*_*k*_ × *P*. Two following ways of standardizing elements of **X**: for an individual *i* in cluster *k*,

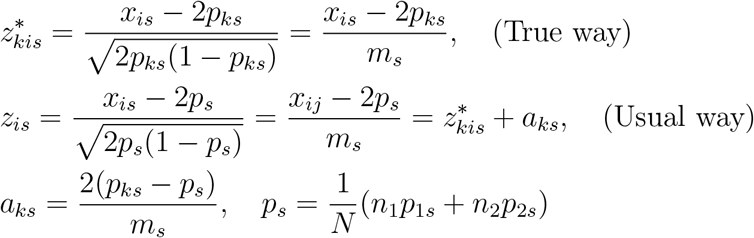

Thus, 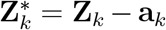 with 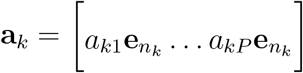. Or, **Z**\* = **Z** − **a** with 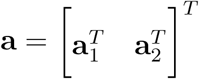.

True GRM and usual GRM can be written as,

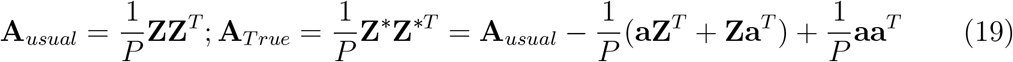

With 2 distinct population subclusters, the first PC of **A**_*usual*_ would be a vector with *v*_1_ for *n*_1_ individuals from cluster 1 and −*v*_2_ for *n*_2_ individuals from cluster 2 with 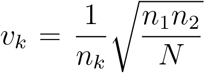 (Patterson et al., 2006; Galinsky et al., 2016). More formally, 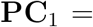 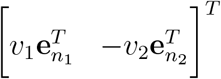. We can write,

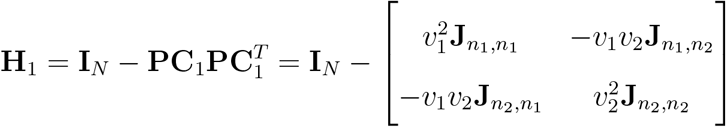

For the *s*-th column of **a** matrix,

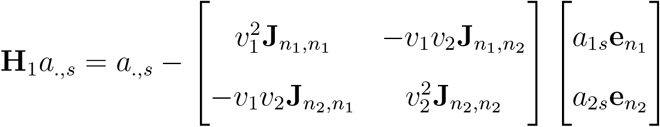

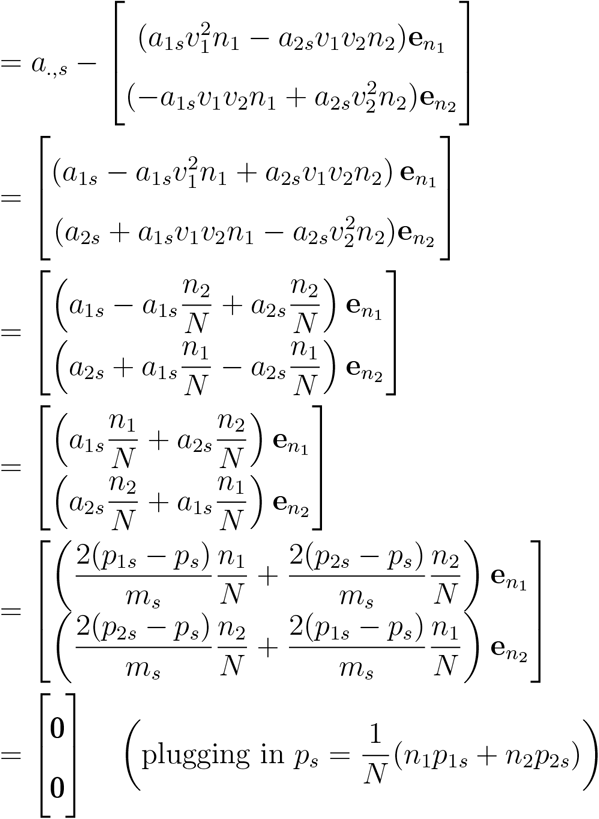

Thus, we find that premultiplying **a** by **H**_1_ gives **H**_1_**a** = **0**. Using this fact in the expressions of equation (2) we get,

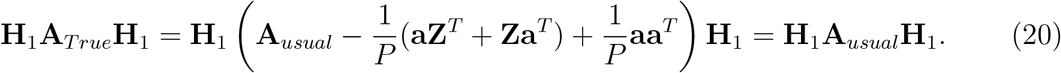

Plugging **H** = **H**_1_ in equation (1) we get,

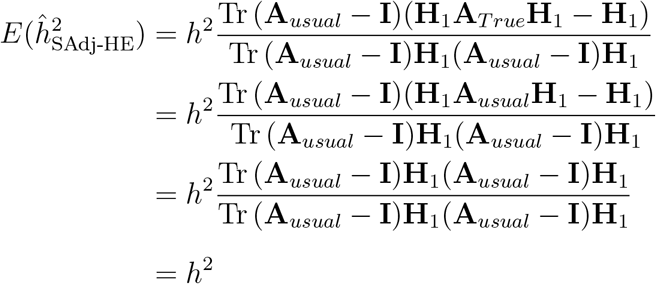

Thus, it shows why Haseman Elston regression with 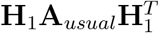 (MMHE) or our proposed Haseman Elston regression with the first PC product adjustment would give us asymptotically unbiased estimate of heritability. When there are more clusters, more PCs would be needed to be considered.

