## Supplemental Tables and Figures for "Estimating SNP heritability in presence of population substructure in biobank-scale datasets"

### Supplementary material

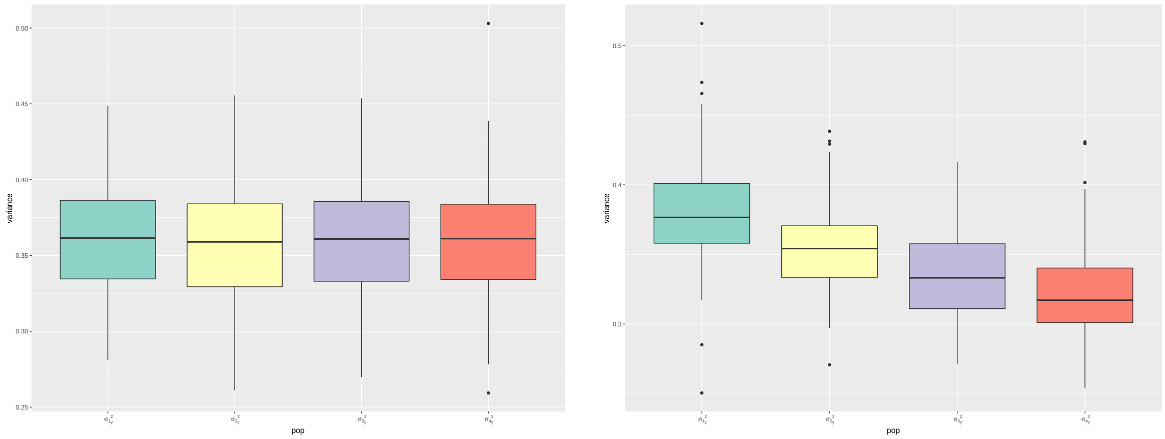


Figure S1. Variance of $\mathbf{y}$ in each sub-population across 100 replicates in simulation 1 with $\theta_k = 0.1$ (left) and simulation 2 (right).

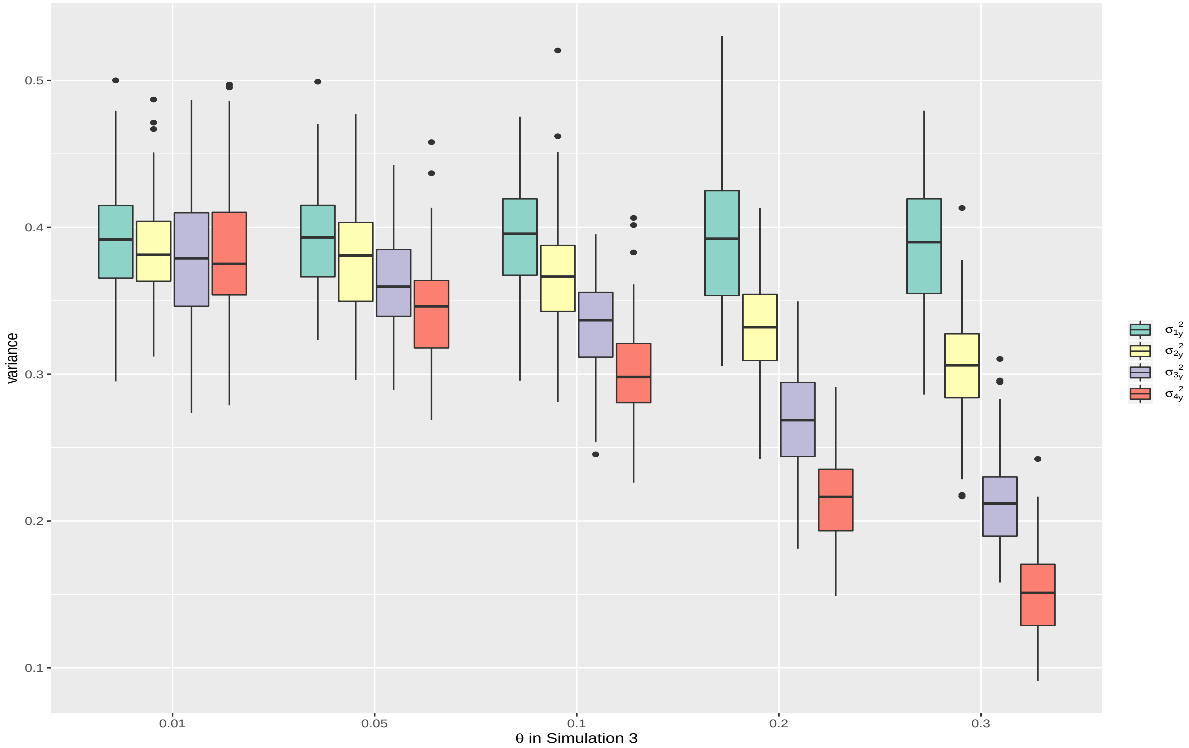


Figure S2. Variance of $\mathbf{y}$ in each sub-population across 100 replicates in simulation 3 with different $\theta_k$

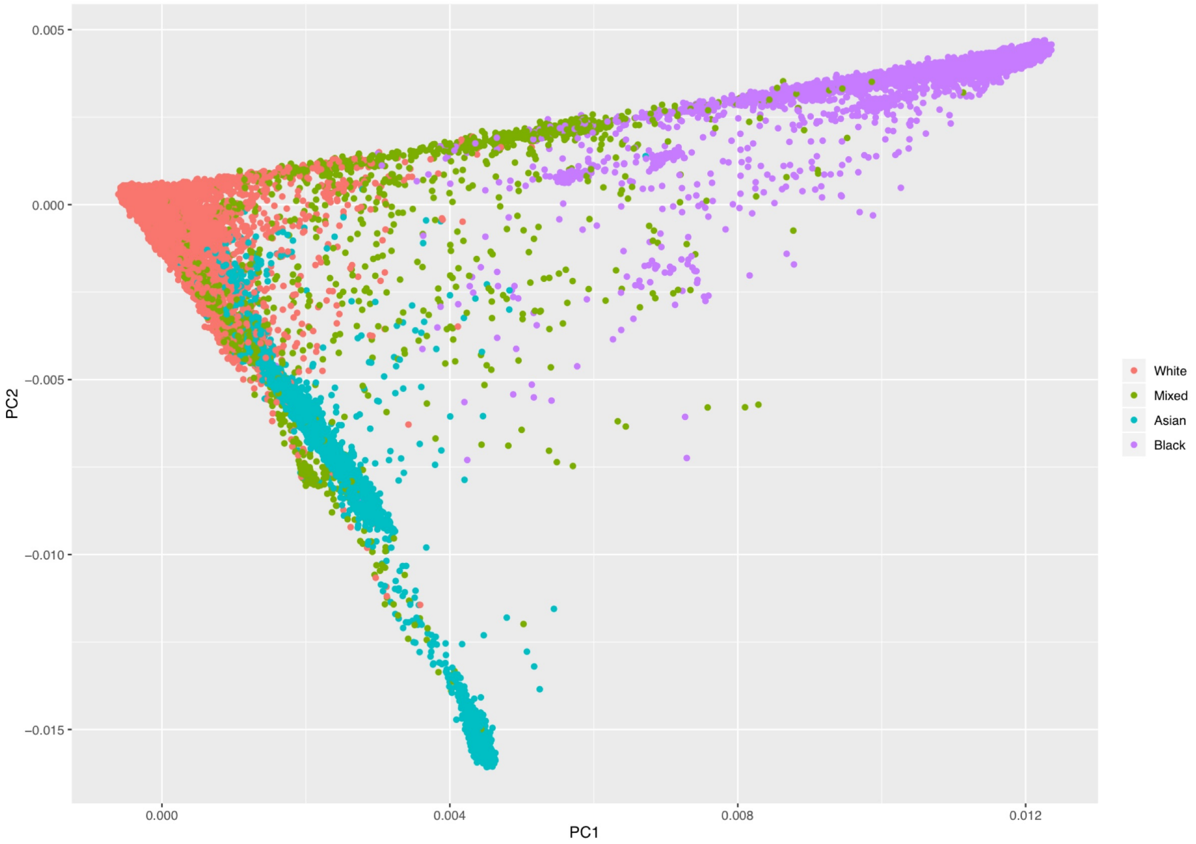


Figure S3. UK Biobank 305k cohort: principal component analysis

Table S1. Empirical Bayes estimates for pairwise Fst in different simulation settings

| Simulation | (1,2) | (1,3) | (1,4) | (2,3) | (2,4) | (3,4) |
|---|---|---|---|---|---|---|
| **Simulation 1** | | | | | | |
| $\theta_k = 0.01$ | 0.005 | 0.005 | 0.005 | 0.005 | 0.005 | 0.005 |
| $\theta_k = 0.1$ | 0.052 | 0.053 | 0.052 | 0.053 | 0.053 | 0.053 |
| **Simulation 2** | 0.039 | 0.054 | 0.067 | 0.067 | 0.081 | 0.096 |
| **Simulation 3** | | | | | | |
| $\theta_k = 0.01$ | 0.003 | 0.005 | 0.007 | 0.003 | 0.005 | 0.003 |
| $\theta_k = 0.05$ | 0.013 | 0.024 | 0.036 | 0.013 | 0.025 | 0.013 |
| $\theta_k = 0.1$ | 0.026 | 0.051 | 0.074 | 0.026 | 0.050 | 0.026 |
| $\theta_k = 0.2$ | 0.055 | 0.103 | 0.144 | 0.054 | 0.102 | 0.053 |
| $\theta_k = 0.3$ | 0.086 | 0.158 | 0.211 | 0.083 | 0.147 | 0.083 |

Table S2. Estimated variability for different traits across ethnicity in 45k sample

| Ethnicity | N | Height | Weight | BMI | DiBP | sysBP | Hip | Waist |
|---|---|---|---|---|---|---|---|---|
| White[1] | 36521 | 85.0 | 255.3 | 23.0 | 116.1 | 390.5 | 83.5 | 184.7 |
| Asian[2] | 5241 | 81.5 | 200.2 | 19.5 | 116.0 | 395.0 | 77.9 | 151.5 |
| Black[3] | 3748 | 73.8 | 241.0 | 27.7 | 125.2 | 374.3 | 101.8 | 148.8 |

[1] Including British, Irish and any other white background.
[2] Including Indian, Pakistani, Bangladeshi, Chinese, Asian British and any other Asian background.
[3] Including Caribbean, African, Black British and any other Black background.

Table S3. Sample mean for different traits across ethnicity in 305k sample

| Ethnicity | N | Height | Weight | BMI | DiBP | sysBP | Hip | Waist |
|---|---|---|---|---|---|---|---|---|
| White[1] | 283718 | 168.9 | 78.2 | 27.3 | 82.1 | 139.7 | 103.4 | 90.2 |
| Mixed[2] | 2151 | 166.9 | 76.4 | 27.3 | 81.3 | 134.0 | 102.8 | 88.6 |
| Asian[3] | 8582 | 163.7 | 71.7 | 26.7 | 82.3 | 136.4 | 100.1 | 89.8 |
| Black[4] | 6026 | 167.4 | 82.3 | 29.4 | 84.7 | 139.7 | 106.0 | 92.8 |

[1] Including British, Irish and any other white background.
[2] Including White and Black Caribbean, White and Black African, White and Asian and any other mixed background.
[3] Including Indian, Pakistani, Bangladeshi, Chinese, Asian British and any other Asian background.
[4] Including Caribbean, African, Black British and any other Black background.

Table S4. Estimated variability for different traits across ethnicity in 305k sample

| Ethnicity | N | Height | Weight | BMI | DiBP | sysBP | Hip | Waist |
|---|---|---|---|---|---|---|---|---|
| White[1] | 283718 | 85.4 | 252.0 | 22.4 | 114.0 | 386.5 | 82.6 | 181.5 |
| Mixed[2] | 2151 | 83.7 | 280.1 | 27.7 | 121.4 | 357.2 | 100.6 | 192.1 |
| Asian[3] | 8582 | 82.1 | 198.6 | 19.3 | 115.9 | 388.2 | 76.2 | 150.8 |
| Black[4] | 6026 | 75.3 | 243.8 | 27.5 | 125.4 | 378.7 | 101.5 | 150.6 |

[1] Including British, Irish and any other white background.
[2] Including White and Black Caribbean, White and Black African, White and Asian and any other mixed background.
[3] Including Indian, Pakistani, Bangladeshi, Chinese, Asian British and any other Asian background.
[4] Including Caribbean, African, Black British and any other Black background.
